# Cryo-EM structure of human somatostatin receptor 2 complex with its agonist somatostatin delineates the ligand binding specificity

**DOI:** 10.1101/2022.01.05.474995

**Authors:** Yunseok Heo, Eojin Yoon, Ye-Eun Jeon, Ji-Hye Yun, Naito Ishimoto, Hyeonuk Woo, Sam-Yong Park, Ji-Joon Song, Weontae Lee

## Abstract

Somatostatin is a peptide hormone regulating endocrine systems through binding to G-protein-coupled somatostatin receptors. somatostatin receptor 2 (SSTR2) is one of the human somatostatin receptors and highly implicated in cancers and neurological disorders. Here, we report the high resolution cryo-EM structure of full-length human SSTR2 bound to the agonist somatostatin (SST-14) complex with inhibitory G (G_i_) proteins. Our structure shows that seven transmembrane helices form a deep pocket for ligand binding and that the highly conserved Trp-Lys motif of SST-14 positions at the bottom of the pocket. Furthermore, our sequence analysis combined with AlphaFold modeled structures of other SSTR isoforms provide how SSTR family proteins specifically interact with their cognate ligands. This work provides the first glimpse into the molecular recognition of somatostatin receptor and crucial resource to develop therapeutics targeting somatostatin receptors.

## Introduction

Somatostatin (SST) is a cyclic peptide hormone composed of 14 amino acids and highly implicated in several disease^1-3^. SST binds to G-protein-coupled somatostatin receptors (SSTRs) inhibiting an adenylyl cyclase via an inhibitory G-proteins^4,5^. SSTRs belong to the class A G-protein-coupled receptors (GPCRs) and there are five isoforms (SSTR1–SSTR5)^6^. Each SSTR isoform is expressed in different tissues and organs and has distinct function^7^. The SST-SSTR axis is highly implicated in several human diseases including acromegaly, cancers and neurological disorders^8-10^. Several SST analogues targeting SSTRs were developed and they are already in clinical use^9^. However, SSTR isoforms have high degree of sequence identity, making it difficult to develop means to modulate each isoform specifically while minimizing off-target effects. SSTR2 is mainly expressed in brain and endocrine tissues and dysregulation of SSTR2 expression is related with neuroendocrine tumors and Alzheimer’s disease^8,11^. In order to understand the molecular and structural basis of somatostatin receptor interaction with its ligand, we determined the cryo-EM structure SSTR bound to its endogenous agonist somatostatin (SST-14) with a complex with inhibitory G-proteins. Our structural work provides insight into the mechanism by which SSTRs recognize its ligand and will serve as a platform to develop selective agonists and therapeutics.

## Results and Discussion

To investigate the molecular basis of the ligand specific binding by SSTR2, we determined the cryo-EM structure of human SSTR2 bound to SST-14 somatostatin peptide. The full-length human SSTR2 in a thermostabilized apocytochrome b562 RIL (BRIL) fusion form and heterotrimeric Gαi1/Gβ1γ2 complex with Gαi1 recognizing scFv16 were separately expressed and purified, and SSTR2-Gαi1/Gβ1γ2-scFv16 complex in the presence of SST-14 cyclic peptide was prepared for structural study (Supplemental Fig. 1). The complex was plunge-frozen and micrographs were collected using Titan Krios 300 keV with a Gatan K2 Summit direct detector in a movie mode. The collected data were processed and refined with Relion 3.1^12^ (Supplemental Fig. 2 and Supplemental Table 1). During the refinement, SSTR2 was separated into two bodies (Body 1: SSTR2+Gαi1β1 and Body 2: Gβ1γ2+scFv16) and two bodies were separately refined. The overall resolution of the whole SSTR2-Gαi1/Gβ1γ2-scFv16 complex is estimated as 3.72 Å at 0.143 criteria of the gold standard FSC, and the resolutions of Body 1 and Body 2 are estimated as 3.65 Å and 3.22 Å respectively. The cryo-EM map was well resolved and the atomic model for the most part of the SSTR2 complex was built on the map (Fig. 1A and Supplemental Fig. 3).

**Figure 1.**
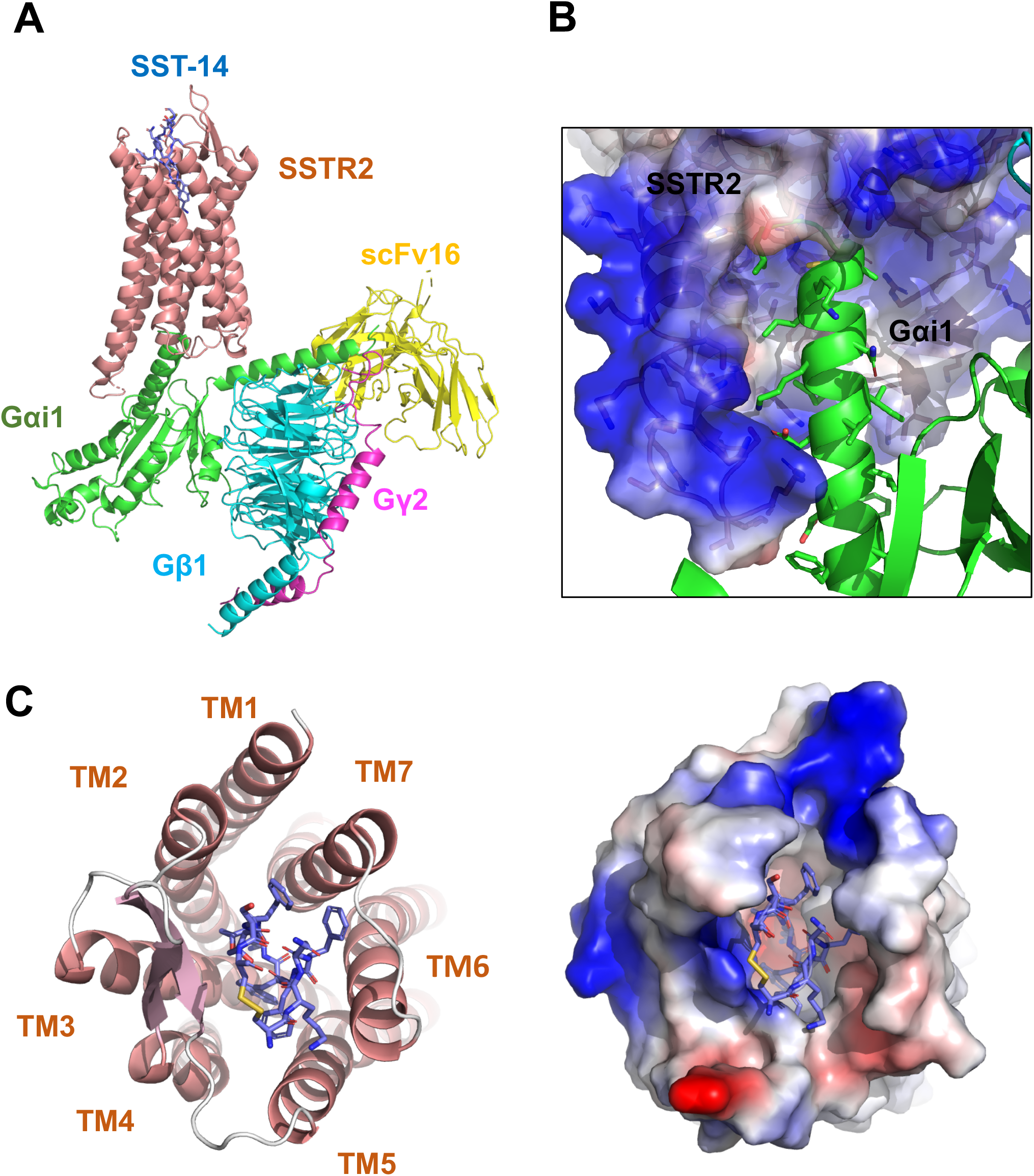
Cryo-EM structure of SSTR2 G-protein complex with SST-14. **(A)** The atomic model of the SSTR2 complex is shown in a ribbon model. SSTR2, Gαi1, Gβ1, Gγ2, and scFv16 are shown in a ribbon diagram and colored in salmon, green, cyan, magenta and yellow respectively. The bound SST-14 cyclic peptide is shown in a ball-and-stick model and colored in navy blue. **(B)** The C-terminal of Gαi1 is inserted in the pocket formed by TM5–7 of SSTR2 forming hydrophobic interactions. **(C)** SST-14 (shown in a ball-and-stick) is bound to the pocket formed by seven TMs of SSTR2 (colored in salmon) at the extracellular side (left panel). SSTR2 is shown in an electrostatic surface representation (right panel).

In the structure, the C-terminal helix of Gαi1 is inserted inside of the seven transmembrane (TM) helices forming hydrophobic interactions with TM 5–7 helices (Fig. 1B), and scFv16 interacts with the N-terminal helix of Gαi1 and a loop protruded from Gβ1, stabilizing the structures. The TM helices of SSTR2 at the extracellular side form a pocket for ligands and SST-14 nestles snugly at the pocket (Fig. 1C). Among 14 amino acids in SST-14, 12 amino acid residues from Cys3 to Cys14 are clearly visible in the cryo-EM map and the positions of the side chains were unambiguously assigned on the map (Fig. 2A). SST-14 is cyclized via a disulfide bond between Cys3 and Cys14 and the region between Phe6 and Phe11 forms a flat sheet (Fig 2B). In this configuration, the disulfide bond is located outward from the ligand binding pocket of SSTR2 and two amino acids of Trp8 and Lys9 is positioned at the bottom of the binding pocket (Fig. 2B).

**Figure 2.**
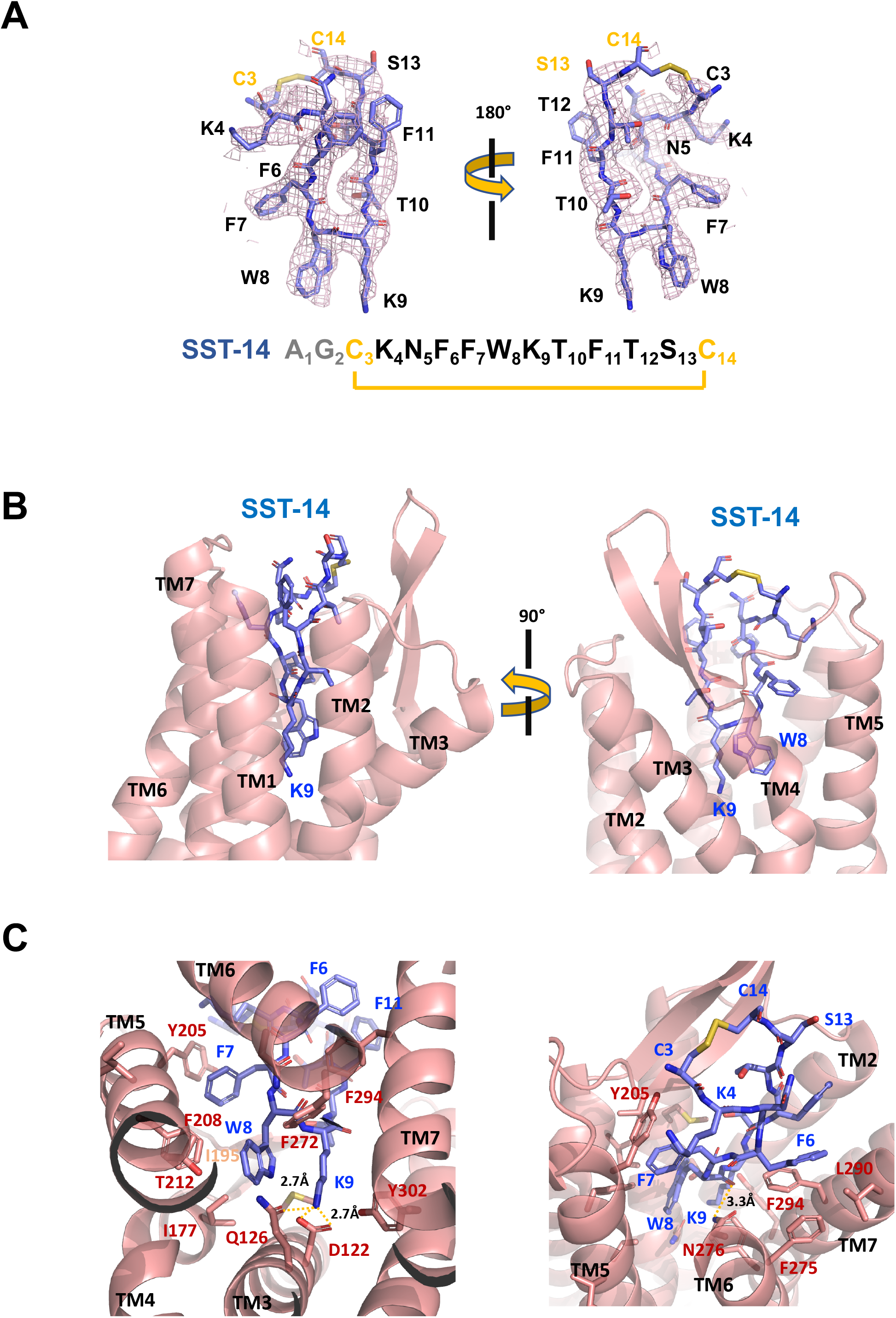
SSTR2 recognition of SST-14 ligand. **(A)** The cryo-EM map near the SST-14 ligand in two different orientations. The sequence of SST-14 is drawn at the below. Two amino acids, which are not visible, are colored in gray. C_3_ and C_14_ makes a covalent bond to form a cyclic peptide. **(B)** The ligand binding pocket of SSTR2 with SST-14 is shown in two different orientations. SSTR2 is shown in a ribbon diagram and SST-14 in a stick model. **(C)** Detailed interactions between SST-14 and SSTR2. The salt bridge between Lys9 and Asp122, and a hydrogen bonding between Lys9 and Gln126 are indicated with yellow dotted lines.

To understand the specific recognition of SST-14 by SSTR2, we examined the detailed interactions between SST-14 and SSTR2. At the bottom of the ligand binding pocket, Trp9 of SST-14 interacts with SSTR2 via a hydrophobic pocket formed by Ile177, Phe208, Thr212 and Phe272 from TM4–6 helices (Fig. 2C). It is of note that Lys9 is the only charged residues among the amino acids of SST-14, which is located inside of the pocket. Lys9 of SST-14 forms a salt bridge with Asp122 with 2.7Å distance and also makes a hydrogen bond with the oxygen atom of Gln126 side chain. In addition, the carbonyl oxygen in the peptide bond between Lys8, and Trp9 form a hydrogen bond with Asn276. While the region of S_13_C_14_C_3_K_4_ is exposed to the solvent, other hydrophobic residues form stable interactions with the hydrophobic residues in the ligand binding pocket (Fig. 2C). Specifically, Phe275, Leu290 and Phe294 residues of SSTR2 accommodate Phe6 residue of SST-14, and Phe7 residue of SST-14 is stacked on Tyr205 coming from the TM5 helix and further stabilized by the interactions with Ile195 and Phe208 residues.

Endogenous agonists for somatostatin receptors are SST-14 and SST-28^4,13^. SST-28 has extra14 residues at N-terminus compared to SST-14 (Fig. 3A). Our cryo-EM structure shows that the region of SST-14 is likely to be sufficient for binding to SSTR2. Consistent with this, SST-28 has a similar binding affinity to SST-14 in a sub-nanomolar range^8^. Besides SST-14, there are several other ligands and drugs that bind to SSTR2 including Cortistatin, Octreotide, Pasireotide, Lanreotide and others. All of these ligands and drugs are cyclic forms of peptides and contain absolutely conserved Trp and Lys residues (Fig. 3A). The interaction analysis based on the cryo-EM structure reveals that these highly conserved Trp and Lys residues tightly interact with SSTR2 via hydrophobic interactions as well as a salt bridge. Therefore, it is likely that the Trp-Lys motif of the ligands and drugs is a major determinant for the binding.

**Figure 3.**
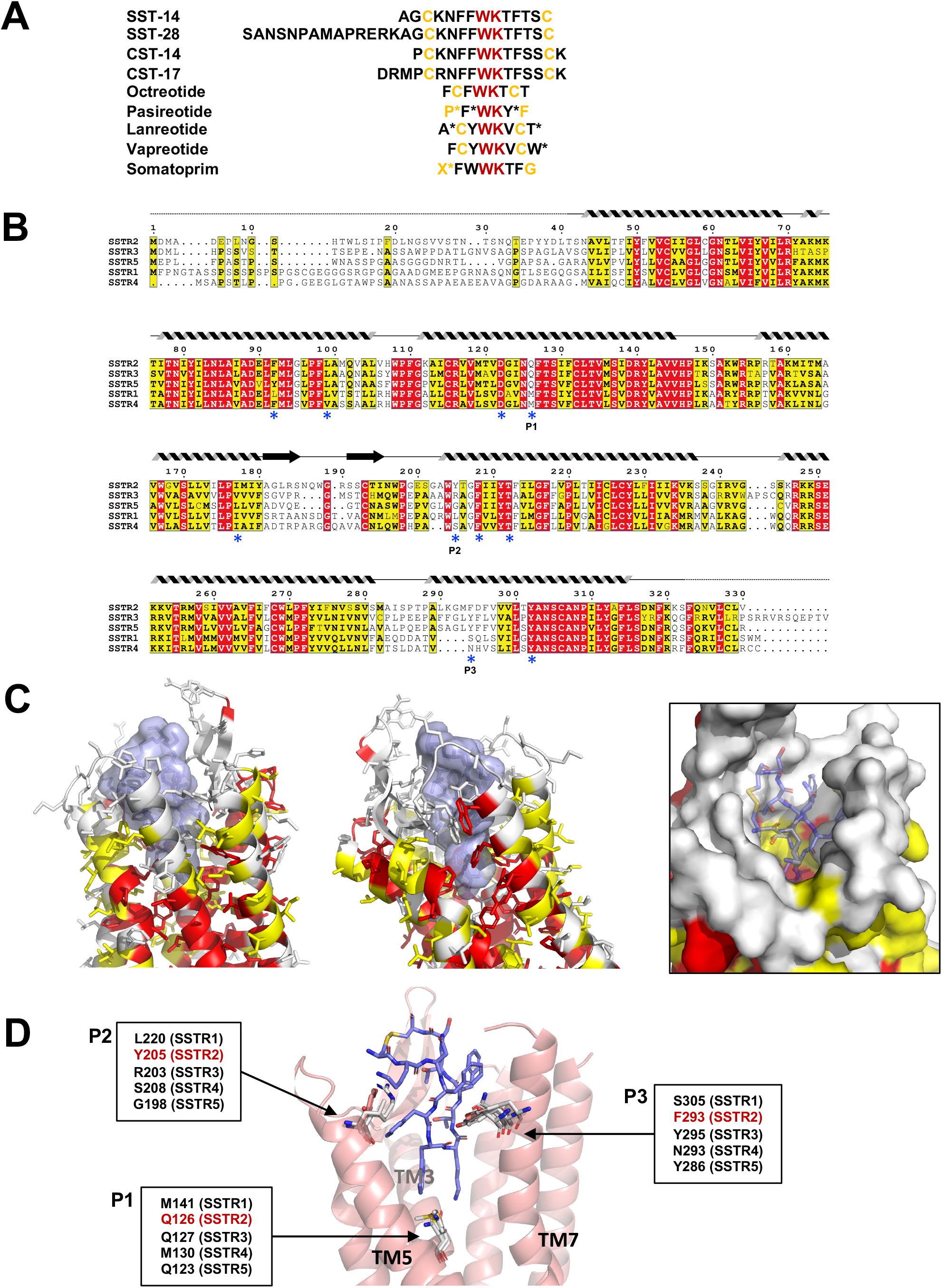
Sequence and structural analysis of ligand binding pockets of SSTR isoforms. **(A)** The sequences of Somatostatins and its analogues. Trp-Lys motif is absolutely conserved among all SSTR binding ligands, emphasizing the importance of the Trp-Lys motif for the function. Modified amino acids are marked with asterisks in Pasireotide (P*: hydroxyproline, F*: 2-phenylglycine, Y*: phenylmethylated tyrosine), Lanreotide (A*: D-2-naphthylalanine, T*: amidated threonine), Vapreotide (W*: amidated tryptophan) and Somatoprim (X*: ω-amino acid). **(B)** A sequence alignment among SSTR isoforms. The amino acids are colored in red or yellow depending the degree of sequence conservation among the isoforms. The secondary structures based on the cryo-EM structure of SSTR2 are shown above the sequences. Critical residues recognizing SST-14 are indicated with asterisks under the sequences. P1, P2 and P3 indicate highly variable sequences among the isoforms. **(C)** The conserved amino acids near the ligand binding pocket are colored according to the sequence alignment in (B). Highly conserved amino acids are clustered near the inner bottom of the pocket, which interacts with the Trp-Lys motif, while amino acids near the upper part of the pocket are not well conserved. **(D)** Three variable positions (P1, P2, P3) in the ligand binding pockets based on the cryo-EM structure of SSTR2 and AlphaFold modeled structure of SSTR1, SSTR3, SSTR4 and SSTR5.

There are five isoforms of somatostatin receptors (SSTR1–5), and the expressed tissues of each isoform and its cognate ligand differs, implying the isoform specific function of somatostatin receptors^8^. To further investigate the ligand specificity of the isoforms, we further examined the ligand binding pocket of SSTRs combining a sequence alignment analysis, our cryo-EM structure of SSTR2 and predicted models of other isoforms from AlphaFold database (https://alphafold.ebi.ac.uk)^14^. To examine the sequence conservation of the ligand binding pocket, we colored the residues near the ligand binding pocket of SSTR2 in red or yellow depending on the degree of conservation based on a sequence alignment among SSTRs (Fig. 3B and 3C). This practice revealed that the residues involved in interacting the Trp-Lys motif are highly conserved. On the other hand, the residues interacting with the other part of SST-14 are highly variable, suggesting that this region contributes to the ligand specificity among SSTR isoforms.

To further understand the binding specificity of SSTR isoforms, we superimposed modeled structures of SSTR1, 3, 4 and 5 isoforms from AlphaFold Structure Database (alphafold.ebi.ac.uk)^14^ on the cryo-EM structure of SSTR2 (Supplemental Fig. 4) and examined the interactions between SST-14 bound to SSTRs. Among the residues in the ligand binding pockets, residues at three positions, which directly interact with SST-14, vary among the isoforms (Fig. 3D). The first position is at Gln126 in SSTR2 located in the bottom of the ligand binding pocket and involved in interacting with the Trp-Lys motif of SST-14. SSTR3 and SSTR5 also has Gln at this position while SSTR1 and SSTR4 have Mets. As this position is located between Trp8 and Lys9, it can be imaged that Gln residue interacts with Lys9 while Met residue interacts with Trp8. The second position is at Tyr205 interacting with Phe7 of SST-14 by stacking interacting between aromatic rings. Each isoform has different residue at this position (Leu220 in SSTR1, Arg203 in SSTR3, Ser208 in SSTR4 and Gly198 in SSTR5). Therefore, it is possible that the residues at this position contribute to the binding specificity of each isoform. The third position is at Phe294 in SSTR2 located on TM7. Phe294 holds Phe6 of SST-14 via hydrophobic interactions. While SSTR3 and SSTR5 have Tyr at this position maintaining the hydrophobicity, SSTR1 and SSTR4 have Ser305 and Asn293 at this position, suggesting that SSTR2, SSTR3 and SSTR5 likely have different binding characteristics compared to SSTR1 and SSTR4. Consistent with this hypothesis, SSTR2, SSTR3 and SSTR5 show nanomolar binding affinity toward Octreotide and Lanreotide drugs while SSTR1 and SSTR4 exhibits micromolar binding affinities ^8,15^. Our structural and sequence analysis reveal subtle differences in the ligand binding pocket in each SSTR isoform, providing a critical clue to understand how each SSTR isoform specifically recognizes its cognate ligand and drug.

In conclusion, our works on the structure of the SSTR2 and SST-14 ligand complex delineate the specific ligand recognition by somatostatin receptors. Furthermore, as somatostatin receptors are highly implicated in several human diseases, our structure work on SSTR2 and SST-14 complex will serve as a fundamental platform to design novel and specific therapeutics to modulate SSTRs.

## Materials and Methods

### Construct design

WT full length human SSTR2 inserted with an N-terminal hemagglutinin (HA) signal sequence, FLAG tag, followed by an 8× His tag and HRV 3C protease cleavage site was cloned into the modified pFastBac1 (Invitrogen, Carlsbad, CA, USA) vector. For the SSTR2 stability, an additional sequence of A1-L106 encoding thermostabilized apocytochrome b562 RIL (BRIL) with the mutations (M7W, H102I, R106L) was added after the 8× His tag at the N-terminus^16^. The heterotrimeric Gαi1/Gβ1γ2 was designed as previously described in mini-Gαq/Gβ1γ2^17^. Single chain antibody scFv16 containing GP67 secretion signal sequence was also inserted into pFastBac1 vector^18^.

### Expression of SSTR2, Gαi/Gβγ heterotrimer and scFv16

SSTR2 and Gαi1/Gβ1γ2 were expressed respectively using Bac-to-Bac Baculovirus Expression system (Invitrogen) in *Spodoptera frugiperda* (Sf9) cells using ESF media (Expression Systems, Davis, CA, USA). Using the high titer virus at a multiplicity of infection (MOI) of 3, Sf9 cells density of 2 × 10^6^ cells/mL in 400 mL biomass were infected. The cells were incubated with shaking at 27 °C for 72 h and harvested, followed by washing with phosphate-buffered saline (PBS), flash-freezing in liquid nitrogen, and storage at −80 °C until further use. The single chain antibody scFv16 was expressed using Bac-to-Bac Expression system in Sf9 cells, and high titer virus was generated. The cells were incubated with shaking at 27 °C for 72 h, and secreted scFv16 in the supernatant was separated from the cells by centrifugation.

### Purification of SSTR2, Gαi/Gβγ heterotrimer and scFv16

SSTR2 frozen pellets were thawed and resuspended at 4 °C with the addition of EDTA-free protease inhibitor cocktail (Sigma Aldrich, St. Louis, MO, USA). The cell membranes were obtained by repeated lysis and dounce homogenization using hypotonic buffer containing 10 mM HEPES (pH 7.5), 10 mM MgCl_2_, 20 mM KCl, and protease inhibitors and hypertonic buffer containing 10 mM HEPES (pH 7.5), 10 mM MgCl_2_, 20 mM KCl, 1.0 M NaCl, and protease inhibitors. Washed membrane fractions were resuspended in buffer containing 30 mM HEPES (pH 7.5), 5 mM MgCl_2_, 10 mM KCl, 500 mM NaCl, 200 μM SST-14, and protease inhibitors. Then, membrane fractions were incubated at 25 °C for 1 h. Then, solubilized in 1% (w/v) n-dodecyl-β-D-maltopyranoside (DDM) (Anatrace, Maumee, OH, USA), 0.2% (w/v) cholesteryl hemisuccinate (CHS) (Anatrace, Maumee, OH, USA) at 4 °C for 3 h. The solubilized solution was isolated by ultracentrifugation at 150,000 × g for 60 min and then supernatant was isolated. TALON IMAC (Clontech) resin was added to the supernatant. The mixture was incubated at 4 °C, overnight. After incubation, the resin-bound SSTR2 was loaded onto a disposable chromatography column (Bio-Rad, Hercules, CA, USA) and the resin was washed with 20 column volumes (CVs) of wash buffer containing 50 mM HEPES (pH 7.5), 500 mM NaCl, 10 mM MgCl_2_, 1% (w/v) DDM, 0.2% CHS (w/v), 5 mM imidazole, 10% (v/v) glycerol, and 50 μM SST-14. Bound proteins were eluted with 10 CVs of elution buffer containing 50 mM HEPES (pH 7.5), 500 mM NaCl, 0.05% (w/v) Lauryl maltose neopentyl glycol (LMNG) (Anatrace, Maumee, OH, USA), 0.005% (w/v) CHS, 300 mM imidazole, 10% (v/v) glycerol, and 100 μM SST-14. PD-10 desalting column (Cytiva) was used to remove the high concentration of imidazole. SSTR2 was then treated overnight at 4 °C with HRV 3C protease. Reverse affinity column was used for the further purification of untagged SSTR2 with buffer containing 50 mM HEPES (pH 7.5), 500 mM NaCl, 0.05% (w/v) LMNG, 0.005% (w/v) CHS, 10% (v/v) glycerol, and 50 μM SST-14. The SSTR2 was collected and concentrated, then loaded onto a Superdex 200 Increase 10/300 GL column (Cytiva) with buffer containing 20 mM HEPES (pH 7.5), 100 mM NaCl, 1 mM MgCl_2_, 0.5 mM TCEP, 0.05% (w/v) LMNG, 0.005% (w/v) CHS, and 50 μM SST-14 via ÄKTA pure system (Cytiva). The fresh SSTR2 was used for SSTR2-Gαi1/Gβ1γ2 complex formation. Gαi1/Gβ1γ2 frozen pellets were thawed and resuspended at 4 °C with the addition of protease inhibitor cocktail. Cells were lysed in lysis buffer containing 20 mM HEPES (pH 7.5), 100 mM NaCl, 1 mM MgCl_2_, 20 mM imidazole, 5 mM β-mercaptoethanol, 100 μM GDP, 1% (v/v) Tergitol-type NP-40 (Sigma), and protease inhibitors. The soluble fraction was isolated by ultracentrifugation at 130,000 × g at 4 °C for 30 min. The Gi heterotrimer containing soluble fraction was purified using Ni-NTA chromatography and eluted with buffer containing 20 mM HEPES (pH 7.5), 100 mM NaCl, 1 mM MgCl_2_, 300 mM imidazole, 5 mM β-mercaptoethanol, and 10 μM GDP. HRV 3C protease was added and the 6× His tag was cleaved at 4 °C for overnight. Reverse affinity column was used for purification of untagged Gαi1/Gβ1γ2. The untagged Gαi1/Gβ1γ2 protein was further purified by size exclusion chromatography (SEC) on a HiLoad 16/600 Superdex 200 column (Cytiva) with following buffer containing 20 mM HEPES (pH 7.5), 100 mM NaCl, 1 mM MgCl_2_, 500 µM TCEP, and 10 µM GDP. The eluted protein was concentrated to 5 mg/mL and stored at −80 °C until further use. The supernatant containing scFv16 was loaded onto HisTrap EXCEL column. The column was washed with 10 CVs of wash buffer containing 20 mM HEPES (pH 7.5), 100 mM NaCl, and 50 mM imidazole. The bound protein was eluted using the same buffer supplemented with 500 mM imidazole. After the eluted protein was concentrated, PD-10 desalting column was used to remove the high concentration of imidazole. C-terminal 6× His tag was cleaved by incubation with HRV 3C protease at 4 °C for overnight. Reverse affinity column was used for purification of untagged scFv16. The scFv16 was further purified by SEC on a HiLoad 16/600 Superdex 200 column (Cytiva) with following buffer containing 20 mM HEPES (pH 7.5) and 100 mM NaCl, Monomeric fractions were pooled, concentrated, and flash-frozen in liquid nitrogen until further use.

### Formation of SSTR2-Gαi1/Gβ1γ2 complex

For forming SSTR2-Gαi1/Gβ1γ2-scFv16 complex, fresh SST-14 bound SSTR2 was mixed with a 1.2 molar excess of Gαi1/Gβ1γ2. The coupling reaction was performed at 25 °C for 1 h and followed by addition of 0.2 unit/mL apyrase (New England Biolabs). After additional 1 h at 25 °C, lambda phosphatase (New England Biolabs) were added. To make SSTR2-Gαi1/Gβ1γ2-scFv16 complex, 1.2 molar excess of scFv16 was added to SSTR2-Gαi1/Gβ1γ2 complex and further incubated at 4 °C for overnight. The SSTR2-Gαi1/Gβ1γ2-scFv16 complex sample in LMNG/CHS was loaded on Superdex 200 Increase 10/300 GL column (Cytiva) equilibrated in buffer containing 20 mM HEPES (pH 7.5), 100 mM NaCl, 1 mM MgCl_2_, 0.5 mM TCEP, 0.001% (w/v) LMNG, 0.0001% (w/v) CHS, and 40 μM SST-14. Peak fractions were concentrated to 2.5 mg/mL for electron microscopy studies.

### Cryo-EM image processing

Cryo-EM data were collected from Titan Krios, Yokohama, Japan. Total 5,523 movies were collected by electron counting mode for 50 frames with a total dose of 55.04 e/Å^2^. Magnification of micrographs was × 75,000, 0.867 Å/pixel. After data collection, image processing was done by Relion 3.1^12^ in a SBGrid package (www.sbgrid.org) ^19^. Initially, collected movies were motion-corrected by MotionCorr2 and ctffind using CTFFIND 4.1 embedded in Relion 3.1. Initial 6,677,042 particles were picked by template-based auto picking. Bad particles were filtered out through several rounds of 2D classification until secondary structures are visible in 2D classes. After 2D classification, selected 2,906,685 particles were subjected to 3D classification with C1 symmetry dividing particles into 8 classes. Among eight 3D classes, three high-resolution classes were selected. Several rounds of 3D classification were done until the final resolution reached 4.08 Å by 3D auto-refine job with final 320,885 particles. The final particles were re-extracted from motion-corrected micrographs with a total dose of 29.24 e^-^/Å^2^, and the resolution was improved up to 3.72 Å (FSC threshold 0.143). For further processing, the whole model was divided into 2 bodies for multibody refinement. Body 1 contains SSTR2+Gαi1/Gβ1+scFv16, and Body 2 contains Gβ1/Gγ2+scFv16. The resolution after multibody refinement and sharpening was 3.65 Å for Body 1 and 3.22 Å for Body 2 (FSC threshold 0.143). The cryo-EM map and the model are to be deposited at EMDB (www.ebi.ac.uk) and RCSB (www.rcsb.org) data base with the accession codes of EMD-32543 and 7WJ5, respectively.

## Acknowledgements

We thank the members of Song and Lee Laboratory for technical assistant and helpful discussion. We also thank Dr. Seung-Hee Lee for critical reading of this manuscript. We thank for the staffs of the cryo-EM facility at RIKEN Center for Biosystems Dynamics Research, Yokohama, for the data collection. This work is supported by a grant (NRF-2020M3A9G7103934 to J.S. and W.L.) from National Research Foundation (NRF) of Korea. The software programs for the processing were supported by SBGrid (www.sbgrid.org). The computing resource was supported by the Global Science Experimental Data Hub Center (GSDC), Korea Institute of Science and Technology Information (KISTI) and by the data analysis hub, Olaf in the Institute of Basic Sciences (IBS) Research Solution Center.

## Author contributions

Y.H., E.Y., J.-H.Y., J.S. and W.L. conceptualized the project. Y.H., Y.-E.J. and J.-H.Y. generated the samples. N.I. and S.-Y.P. collected cryo-EM data, and E.Y. and J.S. processed the data and determined the structure. Y.H., E.Y., J.-H.Y., H.W., J.S. and W.L. analyzed the structure and all authors examined the data. Y.H., E.Y., J.S. and W.L wrote the manuscript. J.S. and W.L. supervised the project and obtained the funds.

