## Supplementary material for "Cryo-EM structure of human somatostatin receptor 2 complex with its agonist somatostatin delineates the ligand binding specificity": Supplmental Figures

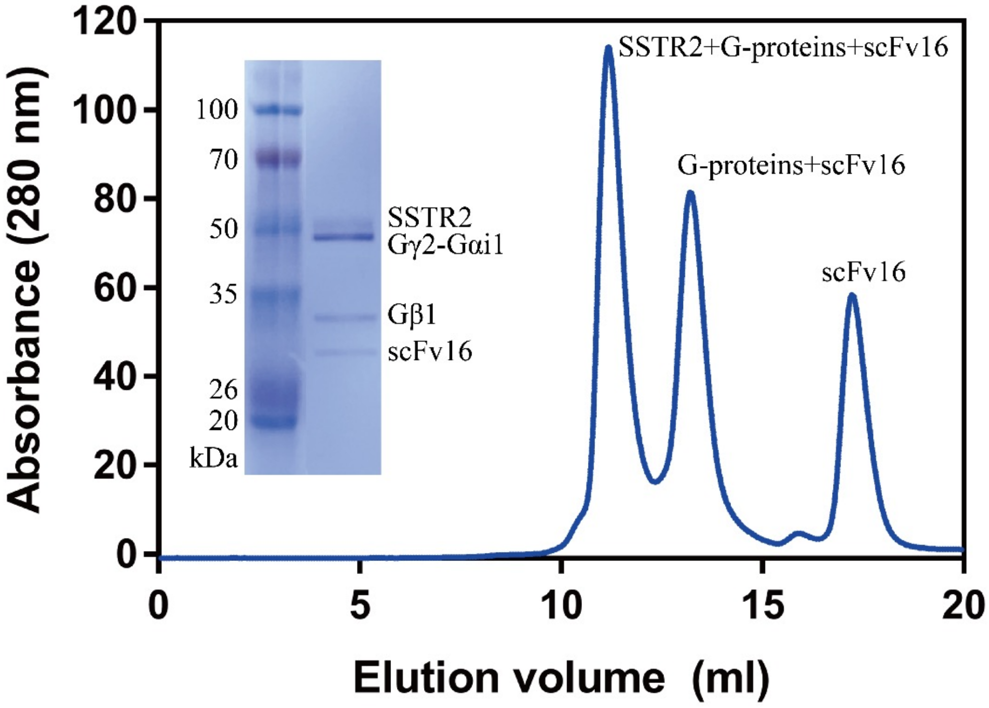


**Supplemental Figure 1. Purification of recombinant human SSTR2-Gαi1/Gβ1γ2-scFv16 complex.**

After injecting the complex sample into a size exclusion chromatography column, three peaks appeared. The complete complex of SSTR2, heterotrimeric Gαi1/Gβ1γ2 and scFv16 was eluted at the first peak around 11.25ml (shown in an SDS-PAGE).

A

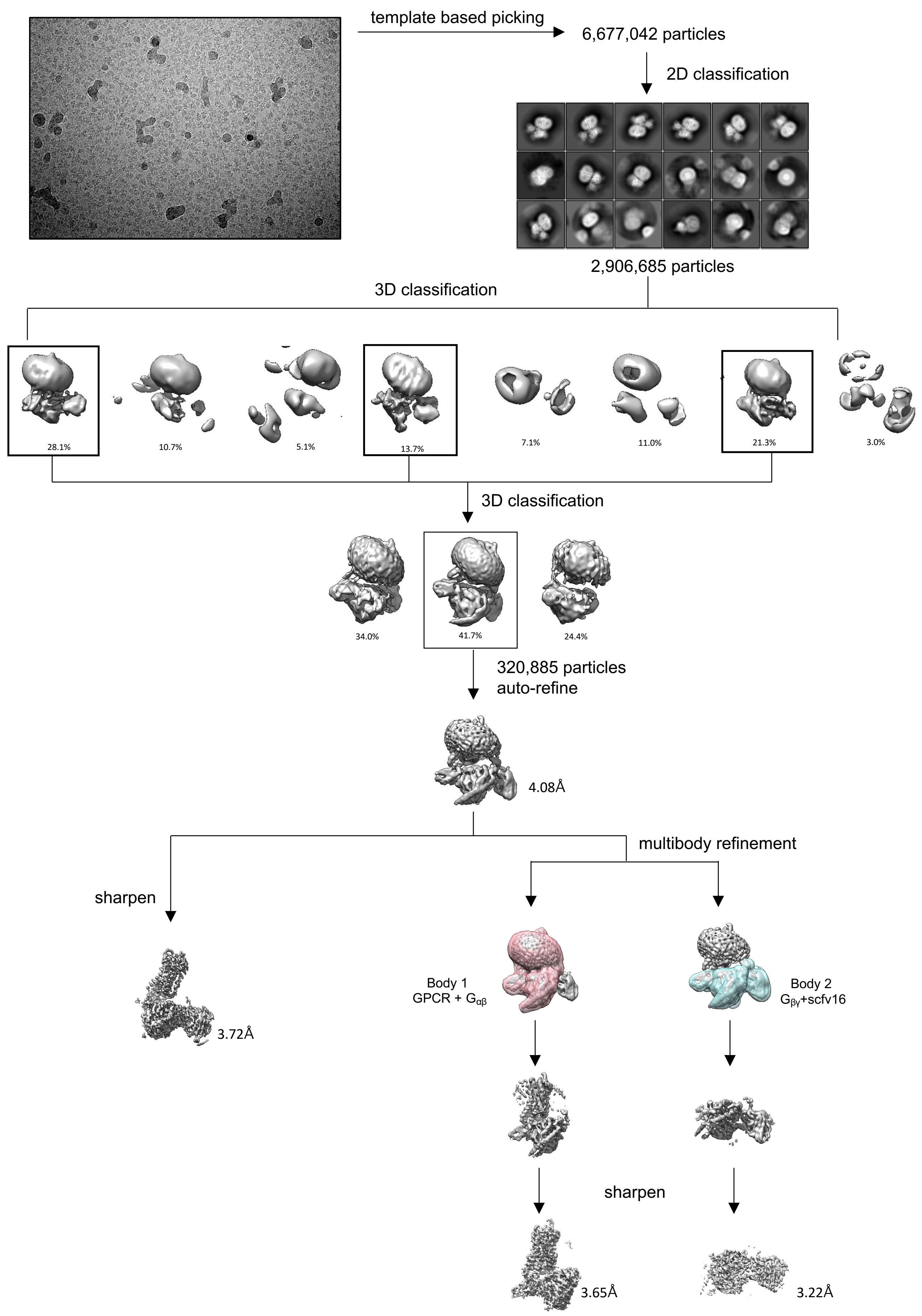
template based picking
6,677,042 particles
2D classification
2,906,685 particles
3D classification
28.1%
10.7%
5.1%
13.7%
7.1%
11.0%
21.3%
3.0%
3D classification
34.0%
41.7%
24.4%
320,885 particles
auto-refine
4.08Å
multibody refinement
sharpen
3.72Å
Body 1
GPCR + $G_{\alpha\beta}$
Body 2
$G_{\beta\gamma}$+scfv16
sharpen
3.65Å
3.22Å

**B**

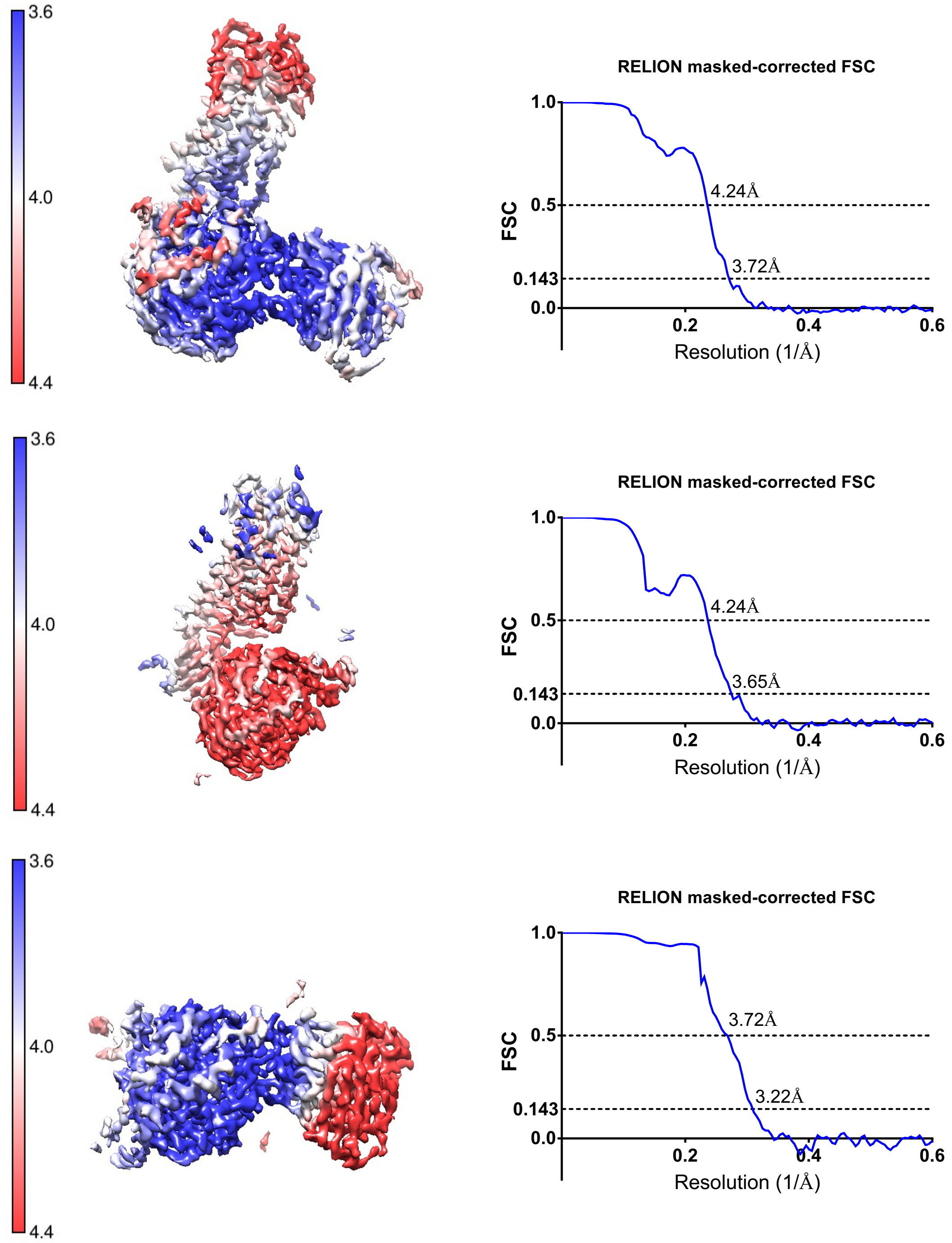


**C**

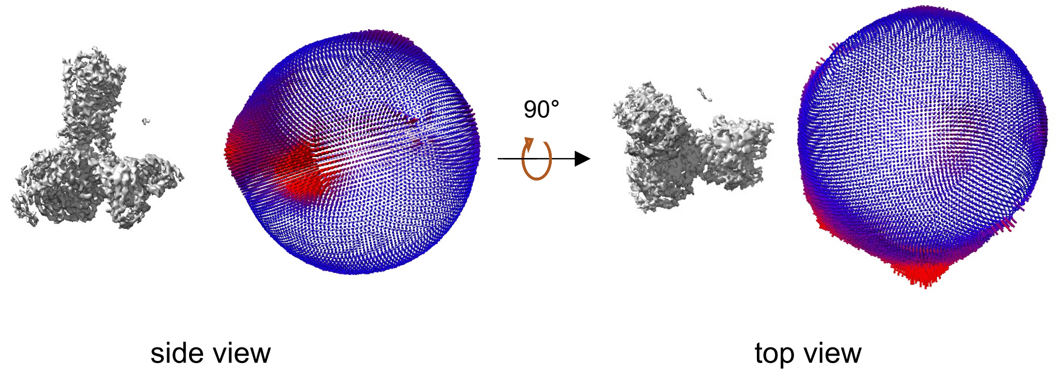


**Supplemental Figure 2. Cryo-EM processing.**
**(A)** The collected data was processed using Relion 3.1. Through 2D and 3D classification, final 320,885 particles were selected for reconstruction. After sharpening, the resolution of SSTR2 complex was determined at 3.72 Å. **(B)** FSC curves of Body 1 and Body 2 as well as complete complex of SSTR2 were obtained. SSTR2 of Body1 showed better resolution than SSTR2 of complete complex, and G-proteins of Body2 showed better resolution than G-proteins of complete complex. **(C)** Euler angle distribution of the particles for the 3D reconstruction.

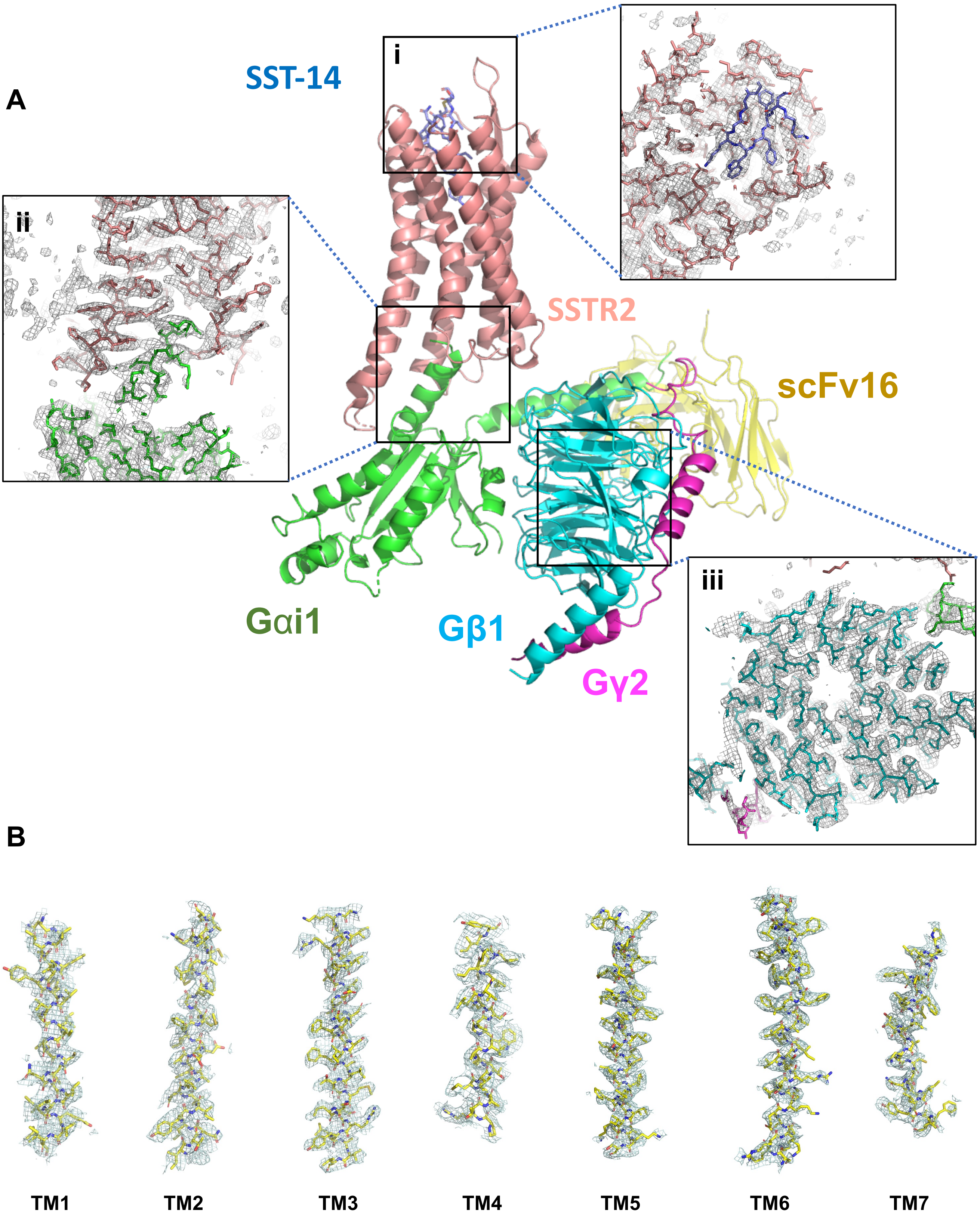


**Supplemental Figure 3. Cryo-EM structure of SSTR2 complex**
**(A)** The cryo-EM maps of SSTR2 complex at the ligand binding pocket (i), the interface between SSTR2 and G-proteins (ii) and the G-protein (Gβ1) (iii).
**(B)** The atomic models of the seven transmembrane helices (TM1: 43–69 a.a., TM2: 77–105 a.a., TM3: 111–145 a.a., TM4: 156–180 a.a., TM5: 203–237 a.a., TM6: 245–281 a.a., and TM7: 288–315 a.a.) are superimposed on the cryo-EM map.

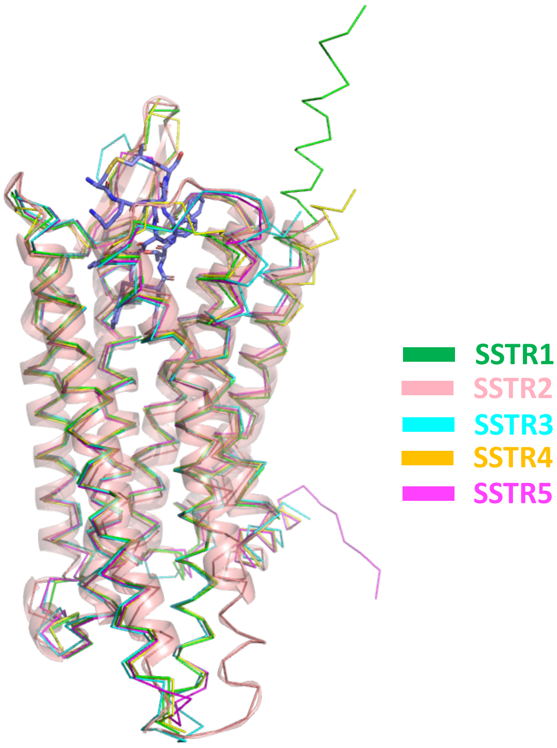


**Supplemental Figure 4. AlphaFold Modeled Structures of SSTR isoforms.**
Structures of SSTR1, 3, 4 and 5 from AlphaFold Protein Structure Database (alphafold.ebi.ac.uk) were superimposed on the cryo-EM structure of SSTR2. SSTR2 is shown in a ribbon diagram in salmon color. SSTR1 is shown as a Cα backbone model and colored in green, SSTR3 in cyan, SSTR4 in gold and SSTR5 in magenta.
