## Supplemental Table1 for "Cryo-EM structure of human somatostatin receptor 2 complex with its agonist somatostatin delineates the ligand binding specificity"

| **Table S1. Refinement Statistics** | | |
|---|---|---|
| **Data Collection and Processing** | | |
| Magnification | | |
| Voltage (kV) | | 300 |
| Total electron exposure / used (e/Å$^2$) | | 55.04 / 29.24 |
| Defocus range (μm) | | -0.8 ~ -2.4 |
| Pixel size (Å$^2$) | | 0.829 |
| Processing program | | Relion 3.1 |
| Obtained / Used micrographs (no.) | | 5,523 / 5,523 |
| Initial / Final particles used (no.) | | 6,677,042 / 320,885 |
| Symmetry imposed | | C1 |
| Resolution (Å) (FSC threshold) | | |
| GPCR + G protein | | 3.72 (0.143) |
| Multibody refinement | GPCR + $G_{\alpha\beta}$ | 3.65 (0.143) |
| | $G_{\beta\gamma}$+scfv16 | 3.22 (0.143) |
| **Refinement** | | |
| Refinement program | | PHENIX |
| Model composition | | |
| Nonhydrogen atoms | | 8,660 |
| Protein residues | | 1,129 |
| R.m.s. Deviation | | |
| Bond Length (Å) | | 0.003 |
| Bond Angle (°) | | 0.609 |
| Validation | | |
| MolProbity Score | | 1.75 |
| Clash Score | | 9.59 |
| Ramachandran Plot | | |
| Favored / Allowed (%) | | 96.31 / 3.51 |
| Outliers (%) | | 0.18 |
| Mask CC | | 0.80 |
